# The origins of time: a systematic review of the neural signatures of temporal prediction in infancy

**DOI:** 10.1101/2025.06.20.659060

**Authors:** Isabelle Rambosson, Damien Benis, Claire Kabdebon, Didier Grandjean, Manuela Filippa

## Abstract

From birth, individuals’ interpersonal dimension is underpinned by progressive learning of social interaction rules, their variations rooted in the temporal prediction of sensory events, and the inferences made about the organization of the social world. How this dimension is progressively structured during infancy and articulated at the neural level is a critical question for cognitive and affective neurosciences. This systematic review aims to define the neural signatures of temporal prediction in newborns and infants and to discuss them in the context of the development of proximal cognitive and affective neural functions. Eight peer-reviewed studies were included, with 228 infants from birth to 9 months of age. Studies have shown that the neural signatures of temporal prediction in infants exhibit a broad cerebral localization, including the anterior and medial parts of the brain, particularly in the frontal and central areas. Temporal prediction mechanisms emerge before birth and range from early sensory-driven responses to more complex top-down processes within the first year of life. While current data do not support a clear longitudinal interpretation, these abilities appear to be shaped by both biological predispositions and experiential factors, with interactive rhythmic and musical activities potentially contributing to their development.

## I. Introduction

In the first months of life, young infants undergo a period of intense brain maturation, during which they begin to construct basic knowledge about their environment and the effects of their actions on it. Bottom-up sensory inputs do not solely drive this learning process, but also rely on top-down mechanisms, as framed by predictive processing models (Emberson, 2017). Predictive coding, as a general framework, provides a compelling explanation for how the brain continuously forms and updates predictions to interact with a dynamic world (Berger & Posner, 2023; Köster et al., 2020).

Prediction, in its temporal articulation, lies at the intersection of theoretical models, methodological advances, and emerging research in infants. This review examines the neural underpinnings of temporal prediction in newborns and infants, underlying the current lack of knowledge regarding the organization of these brain mechanisms early in life.

### 1.1 Theoretical framework

#### The predictive brain in adults and infants

Among the theoretical models that have shaped our understanding of infants’ cognitive development, predictive coding has emerged as a particularly influential framework, integrating neurodevelopmental, computational, and probabilistic principles. The predictive coding theory (Friston, 2005; Rao & Ballard, 1999) is a unifying brain neuro-computational conceptualization, based on Bayesian inference (Perfors et al., 2011), in which the organism builds probabilistic models, by acquiring statistical regularities from the environment (Clark, 2013) and constructing internal generative models of the world (Friston, 2010, 2012; Wolpert et al., 1995). There is ample evidence that the adult brain is highly predictive and follows the principles of the predictive coding theory (see Alexander & Brown, 2019, for example; Berger & Posner, 2023, for a review; Carlsson et al., 2000; Kastner et al., 1999; Zhang et al., 2019).

Recent research suggests that a predictive-processing framework, as a broader reference frame, may also provide a unifying perspective for understanding young infants’ cognitive processes (see Köster et al., 2020, for a review), covering an array of fundamental knowledge about their social and physical environment, dynamically coordinating in time. Indeed, in the early phases of development, the brain must improve inferences based on sensory inputs – by top-down or feedback neural signals – and minimize prediction errors – through bottom-up processes (Baek et al., 2020). Based on this prediction error related to imprecise or false predictions, the model from which predictions were generated is updated (Berger & Posner, 2023).

The above-mentioned mechanisms, linking past and future inputs, enable our cognitive and affective processes to engage with, react to, and adapt to a constantly changing environment (Baek et al., 2020; Berger & Posner, 2023).

#### Statistical and associative learning as a function and a result of the predictive brain

Complementary to predictive coding, statistical and associative learning provide key mechanisms through which infants acquire structured knowledge about their environment. Statistical learning can be defined as the ability to implicitly extract and detect recurrent patterns or regularities in sensory inputs – such as in melodies, speech, or visual scenes – without the need for explicit instruction (Aslin & Newport, 2012; Bogaerts et al., 2023; Saffran, 2024). As a core cognitive process embedded within the predictive processing framework, it enables the anticipation of future events by leveraging probabilistic regularities in the environment (Daikoku & Yumoto, 2023; Wang et al., 2025).

During development, infants systematically use statistical learning abilities, for example, to learn about auditory (i.e., linguistics; Saffran, 2020), visual, and proprioceptive inputs (Trainor, 2012). In parallel, associative learning supports the formation of links between distinct stimuli – ranging from concrete sensory events to abstract concepts (Abd El-Hay, 2019; Lafontaine et al., 2020; Mason et al., 2024), through mechanisms shaped by prediction errors (Holland & Schiffino, 2016). Developmental psychologists often capitalized on associative learning tasks to address general questions about infants’ cognition and learning (Albareda-Castellot et al., 2011; Hochmann et al., 2011; Kovács & Mehler, 2009a, 2009b; McMurray & Aslin, 2004).

#### Temporal structure and predictions

A key domain where predictive processing unfolds is the temporal structure of sensory input, which enables the brain to anticipate when events are likely to occur. Temporal prediction, or the ability to assess the short-term evolution of forthcoming events in time (Tomasi et al., 2015), relies mostly on implicit basic temporal processing mechanisms such as temporal processing and timing prediction.

The temporal structure of the stimuli in the environment plays a crucial role in infants’ predictive processes, and it constitutes the core element of the present review. Here, we specifically focus on the temporal dimension as the organizing principle of interpersonal coordination. This focus stems from increasing evidence that timing is not merely a supporting element of interaction, but a primary framework through which infants learn to anticipate and respond to the social world.

Indeed, from birth, infants need to extrapolate regularities from the sensory world and coordinate their sensory and motor experience with the physical and human environment (Mento & Granziol, 2020). This coordination process is mainly based on the temporal dimension (Mento & Granziol, 2020). The development of an accurate temporal structure of the inputs/outcomes is critical for the formation of expectations (Mento & Granziol, 2020) in interpersonal synchrony, such as early dyadic and triadic exchanges.

Synchrony, or the temporal coordination of micro-level social behaviors, is based on shared and repetitive rhythmic sequences (Bernieri et al., 1988; Feldman, 2007b). Interindividual synchrony relies on anticipating each other’s actions to coordinate in time (Morganti et al., 2008; Sebanz & Knoblich, 2009). Mothers and their infants are involved in interpersonal synchronization from a very early age (Condon & Sander, 1974; Jaffe et al., 2001). During their 3^rd^ month of life, infants start to actively engage in face-to-face interactions, centered on the synchrony of vocalizations, non-verbal cues, gaze, and shared-emotion expressions (Stern, 1985; Tronick, 1989). Interaction synchrony is fundamental to infants’ cognitive, social, and emotional development (Feldman, 2007a; Jaffe et al., 2001) and relies on anticipatory abilities, which are crucial for the prediction of others’ actions and for producing a coordinated response in time (Crown et al., 2002; Jaffe et al., 2001). A good example of these phenomena is the well-known “peek-a-boo” game in which an adult and an infant engage in violating and confirming time expectations, underpinning infants’ time prediction abilities (Baek et al., 2020).

### 1.2 Methodological approaches used to study prediction

#### Deviance detection and expectancy violation in adults (ERPs and oscillatory dynamics)

In adults, predictive coding, in particular in the auditory domain, is supported by deviance detection. The latter can be decomposed into two main mechanisms (i.e., prediction error and repetition suppression; see Carbajal & Malmierca, 2018, for a review). Deviance detection, which is an automatic response to a divergence (such as a violation) in a stimulation previously processed, allows, in the predictive coding framework, the updating of the prior representational model and favors the integration of this novel information within the previously established framework (Winkler & Schröger, 2015).

In the auditory domain, deviance detection is commonly studied using the auditory oddball paradigm, which consists of introducing a new sound after a series of repeated sounds. Such a sudden shift typically elicits an event-related potential (ERP) called the mismatch response (MMR) or mismatch negativity, characterized by a frontal positivity synchronized with a posterior negativity occurring 200-400 ms after the deviant stimulus (Basirat et al., 2014; Friederici et al., 2002). Subsequently, this early response is frequently followed 700 ms later by a late frontal Negative Slow Wave (NSW; Basirat et al., 2014; Dehaene-Lambertz & Dehaene, 1994). This ERP response has been hypothesized to originate from neuronal populations over a network comprised of the auditory midbrain and thalamus, displaying stimulus-specific adaptation (SSA, i.e., a modulation of firing rate in response to a deviant stimulus compared to a series of similar stimuli); triggering in this population a reduction over time of the firing response (Carbajal & Malmierca, 2018).

At the same time, considering the oscillatory dynamics related to expectancy violation, a theta synchrony increase has also been observed in a prefrontal subthalamic network in response to error monitoring and expectancy violation (Zavala et al., 2018), and a theta-gamma coupling has been linked to predictive coding in a syllable representation paradigm (Hovsepyan et al., 2020) in adults. In adults, theta band activity has also been shown to consistently decrease along with the learning of a repeated sequence in line with SMA predictions (Crivelli-Decker et al., 2018).

#### Deviance detection and expectancy violation in infants (ERPs and oscillatory dynamics)

Neural mechanisms involved in deviance detection and expectancy violation are already present in newborns (Alho et al., 1990; Bisiacchi et al., 2009; Cheour-Luhtanen et al., 1995; Cheour et al., 1998; Mahmoudzadeh et al., 2017; Mento et al., 2010), as demonstrated by the elicitation of mismatch negativity (MMN) through oddball paradigms, both in newborns (Cheour et al., 2002; Winkler et al., 2003) and during the fetal period (Draganova et al., 2005; Draganova et al., 2007). As shown by Basirat et al. (2014), 3-month-old infants are already able to extract temporal violations, dissociating local – when the stimulus expectancy is induced by a narrow transitional probability – from global – in the presence of non-adjacent, higher-level rules that supersede local transitions – violations in an auditory paradigm. Infants process these violations *via* two partially distinct brain systems (i.e., MRR/late frontal negative response for the local deviance condition *vs.* a shorter early mismatch response and a longer late response with cortical sources including the left inferior frontal region in the global deviance condition) (Basirat et al., 2014).

Considering the oscillatory dynamics, findings by Köster et al. (2019, 2021) have shown that unexpected (in contrast to expected) events were linked to theta brain activity in 9-month-old infants, as a key element in the processing of prediction errors, over an extended time window (2 sec) and all the electrodes recorder on the scalp (based on 30 electrodes; Köster et al., 2021).

Findings of theta modulation in infants, as well as accurate interval timing and beat processing within one week of birth in premature (32 weeks gestational age or wGA) and full-term babies (39 wGA) (Edalati et al., 2023; Háden et al., 2024), therefore suggest that adult-like processes are already in place early in human development.

In sum, decades of research show that newborns have neural mechanisms involved in temporal expectancy violations, with the auditory oddball paradigm used as an effective tool for studying these processes. Recent research has highlighted the importance of theta brain activity in processing prediction errors, emphasizing the complexity and persistence of infants’ responses to unexpected events.

#### Associative learning and anticipatory paradigms

In these experiments, infants are typically taught an association between the presentation of exemplars of a specific auditory or visual (or abstract) category and the subsequent appearance of a reinforcer on the left or right side of the screen. After a familiarization phase, infants (aged from 6 to 12 months) are presented with novel exemplars of the same category, and their ability to correctly anticipate the appearance of the reinforcer on the left or right side of the screen is assessed as a marker for their categorization abilities (see Albareda-Castellot et al., 2011, for example; Hochmann et al., 2011; Kovács & Mehler, 2009a, 2009b; McMurray & Aslin, 2004).

### 1.3 Relevant literature

#### Behavioral and physiological studies on (temporal) prediction Prediction

Predictive abilities have been investigated behaviorally in the visual modality in infancy by monitoring looking time after habituation, showing that infants as young as newborns, as well as 2-month-old infants, can extract statistical regularities (Bulf et al., 2011; Kirkham et al., 2002). In addition, by tracking anticipatory eye movements, some studies have demonstrated that, from 3 months of age, infants’ brains can predict the position of a moving object (Canfield & Haith, 1991; Gredebäck & von Hofsten, 2004; Gredebäck et al., 2002) or the end state of a motor action (Ambrosini et al., 2013; Cannon & Woodward, 2012; Falck-Ytter et al., 2006; Kanakogi & Itakura, 2011). Using probabilistic inference in an eye-tracking paradigm, recent work additionally demonstrated that when a situation is uncertain, the brain of 12-month-old infants can spontaneously anticipate the outcome of a scene, demonstrating untrained proactive anticipatory behavior in young infants (Téglás & Bonatti, 2016).

#### Temporal prediction

At a behavioral level, temporal predictions have been investigated in infants with interesting results. In one study, where 6- and 12-month-old infants’ eye movements were recorded while observing rational and non-rational feeding actions, Gredebäck and Melinder (2010) reported a change in pupil size – signaling surprise – during violations of infants’ expectations concerning rationality (i.e., in non-rational feeding actions). In another study, Gredebäck et al. (2018) demonstrated that the ability to predict actions in 6-month-old infants is associated with the inclination to react with surprise when social interactions deviate from what was expected. In the auditory modality, Werner et al. (2009) demonstrated using a fixed presentation rate of tone (i.e., fixed condition) and a mixed condition that infants (aged from 6 to 9 months) better detect a tone when it occurred either before or at an expected time following a cue; showing that infants demonstrate greater sensitivity to sounds that align with their temporal predictions.

#### Brain studies on prediction

“Violation”, “prediction”, “expectation”, and “anticipation” are related constructs that reflect a continuum from basic detection to more complex representational and preparatory processes. These terms are often used interchangeably in the literature, creating conceptual ambiguity (Bubic et al., 2010). To clarify this terminology, a violation can be defined as a mismatch between predicted and actual sensory inputs (García Alanis et al., 2023); prediction as a model-based process of forecasting forthcoming inputs (Friston, 2005, 2010); expectation as a representation, shaped by prior experience, that encodes the most likely outcomes (Bubic et al., 2010); and anticipation as a preparatory or readiness state oriented toward those expected outcomes (Andrzejewski et al., 2019).

Together, these definitions describe a dynamic predictive framework in which cognitive and neural systems continuously generate, refine, and act upon internal models of future events.

Despite this confusion in terminology, the idea of brain mechanisms dedicated to anticipation and prediction has been around for decades, as shown by Piaget’s quotations below:

> “*Anticipatory function […] is to be found over and over again, at every level of the cognitive mechanisms and at the very heart of the most elementary habits, even of perception.*”
>
> (Piaget, 1971, p. 191).
>
> “*[…] The function of anticipation is common to cognitive mechanisms at all levels.*”
>
> (Piaget, 1971, p. 193).

If in newborns, dedicated neural mechanisms involved in expectancy violation are already in place (Mahmoudzadeh et al., 2017), it has been much less investigated whether and how early the ability to use this implicit information can be translated into specific, cortical anticipatory processes for future events.

Infants can predict, using, e.g., EEG signals as measures, implying that predictions are inferred from implicit measures, not only the presentation of sequences of stimuli but also the consequences of their actions in the environment (i.e., behavioral inferences). From the age of 2 months, infants consolidate an understanding of the causal link between one of their actions (a foot movement) and the movement of an object linked to it. At 9 months of age, the infant’s brain can, therefore, predict the recurrence of this causal link (Kayhan et al., 2019). Moreover, if the prediction is disregarded (e.g., the object is detached from the foot and no longer moves), infants tend to increase their motor actions to reproduce the coupled event, thereby adapting the predictive model accordingly (Kayhan et al., 2019; Wen & Imamizu, 2022). Interestingly, infants predict an action better when they can perform it, reinforcing the link between action production and perception (Keysers & Perrett, 2004).

Moreover, over the past decade, some research has attempted to identify neural correlates of predictive processing in infancy using associative learning tasks. In 12-month-old infants, Kouider et al. (2015) showed that the acquisition of arbitrary audio-visual associations could modulate visual brain responses, leading to enhanced EEG activity during early processing stages for expected events and increased neural activity during late processing stages for unexpected visual events. Subsequent research using similar associative learning tasks also found correlates of increased neural processing in response to expected events in 5-month-old infants (Kabdebon & Dehaene-Lambertz, 2019) and a late difference between expected and unexpected audio-visual associations in 5- to 6-month-old infants (Kabdebon & Dehaene-Lambertz, 2019; Rohlf et al., 2017). Neural signatures of audio-visual associations have also been investigated (using EEG) in later developmental stages in the context of early word learning using bimodal object-word priming designs. These studies report that consistent object-word priming typically elicits an increase in a frontally distributed N200-500 component and a decrease in the centroparietal N400 component, in 6- to 9-month-old infant ERPs to the word onset (Friedrich & Friederici, 2011; Friedrich et al., 2015, 2017), which could be interpreted as cross-modal/semantic prediction effects. Using functional near-infrared spectroscopy (fNIRS), Emberson et al. (2015) presented 6-month-old infants with a systematic audio-visual association, and they were able to record brain activations over the occipital cortex even when the image was unexpectedly omitted after the auditory cue, similar to the response to the actual image presentation. Notably, this occipital activation was not recorded when the auditory cue did not predict any visual event. These results are suggestive of expectation-based feedback signaling in infancy, although atypical in premature (< 33 wGA) babies (measurements have been realized before infants exhibited clinically identifiable developmental delays at 6 months of corrected age; Emberson, Boldin, et al., 2017), compatible with the predictive coding framework. Finally, a few additional EEG studies from 4 months of age investigated anticipatory pre-stimulus activity in similar audio-visual associative learning tasks in infants, reporting the build-up of a centrally distributed component during the delay between the predictive auditory cue and the onset of the visual stimulus (Kabdebon & Dehaene-Lambertz, 2019; Mento et al., 2022; Mento & Valenza, 2016) comparable to the Contingent Negative Variation (CNV), a neural signature of expectancy-based cortical activity, observed in adult participants.

### 1.4 Adopted perspective and aims of the present review

The adopted perspective in this review suggests that complex predictive abilities emerge early in infants and newborns. Although the basic ability to detect environmental regularities is evident from early in life – partly due to genetic influences and already observable in newborns’ sensory processing and brain network organization – the neural systems that support more sophisticated, prediction-based anticipatory behaviors evolve in complex, nonlinear ways across development. This evolution is shaped by the dynamic interaction between ongoing neurobiological maturation (including processes like myelination, axonal development, and the formation of long-range neural pathways), individual experience, and inherited factors (Mento & Bisiacchi, 2012). Together, these elements support the progressive refinement, specialization, and robustness of brain function as it transitions toward a mature, adult-like architecture.

To date, comprehensive reviews addressing the emergence and organization of the neural correlates of temporal prediction in infants are lacking.

Therefore, in the present review, we aim to disentangle the neural networks involved in temporal predictive abilities across infancy. It is important to note that all studies on infants’ anticipation and prediction imply the temporal dimension; however, we specifically aimed to focus only on studies where the temporal structure (or dynamic unfolding) of the stimuli is manipulated, and thus where the temporal dimension of the stimuli, rather than the stimuli’s characteristics, varied.

The delineation of the defining characteristics and underlying neural substrates of temporal prediction phenomena holds promise for advancing our understanding of the genesis and maturation of core cognitive functions intricately linked to dynamic temporal processes. In the long term, the potential broader impacts of our investigation stand to significantly contribute to the identification of deviations in these capacities at an early age.

## II. Methods

The search and selection strategies have been conducted following the PRISMA (Preferred Reporting Items for Systematic Reviews and Meta-Analyses) 2020 guidelines (Page et al., 2021).

### 2.1. Literature search strategy

A web-based search strategy was elaborated to identify articles related to the neural signatures of temporal prediction in infancy, including infants from birth to 12 months. To conduct a thorough web-based article search, articles were searched in two electronic databases: CINAHL and Web of Science.

The search was conducted using the keywords mentioned below (Box 1). To find the most plausible possibilities regarding the investigated subject, the terms *Infant* and *Brain* remained fixed in all searches.

Box 1. Search terms

Anticipation, auditory anticipation, auditory violation, baby, brain, cortical, infant, neonate, neural, newborn, prediction, predictive coding, rhythmic violation, sensory prediction, and top-down.

### 2.2. Inclusion and exclusion criteria for the selection of eligible studies

The criteria for inclusion of studies were as follows: (a) published in English, (b) empirical peer-reviewed studies embracing a clear methodological stance, (c) on temporal prediction, (d) 0 to 12-month-old infants, and (e) with brain outcomes. Multiple findings of the same article were treated as a single study. Books, book chapters, discussions, theoretical papers, reviews, theses, and dissertations were excluded.

#### Excluded studies on infants’ prediction: which and why?

The present systematic review aimed to highlight the neural signatures of temporal prediction in infancy. Therefore, we excluded all the papers that did not contain an explicit/strict violation of the temporal structure of a series of regular stimuli. Thus, we excluded the studies employing oddball paradigms that are not based on a rhythmical violation, in which the infant perceives a difference between what is currently presented and what was presented before (e.g., Ackles & Cook, 1998), which translates into terms of deviance or novelty effect during development (Kushnerenko et al., 2013).

In the same vein, we also excluded all the paradigms based on the violation of two paired stimuli (e.g., audio and visual stimuli; see Kersey & Emberson, 2017; Rohlf et al., 2017; Wang et al., 2022, for examples). We excluded studies simply changing the order of the elements in a sequence (e.g., Basirat et al., 2014), or the order of stimulus presentation (e.g., Baek et al., 2022), and studies with repetition suppression paradigms (Emberson et al., 2019; Emberson, Cannon, et al., 2017). In general, it follows that all the studies investigating associative learning paradigms, where the association between two or more stimuli was violated, were excluded.

### 2.3. Results of the search

4135 potentially relevant references were found in the primary literature search. 426 articles were excluded based solely on the title or abstract. We identified 124 articles for a relevant full-text review. Based on a critical review (using the criteria mentioned in the previous paragraph) of the full text, only 8 articles were identified as eligible for inclusion (cf. Figure 1).

**Figure 1.**
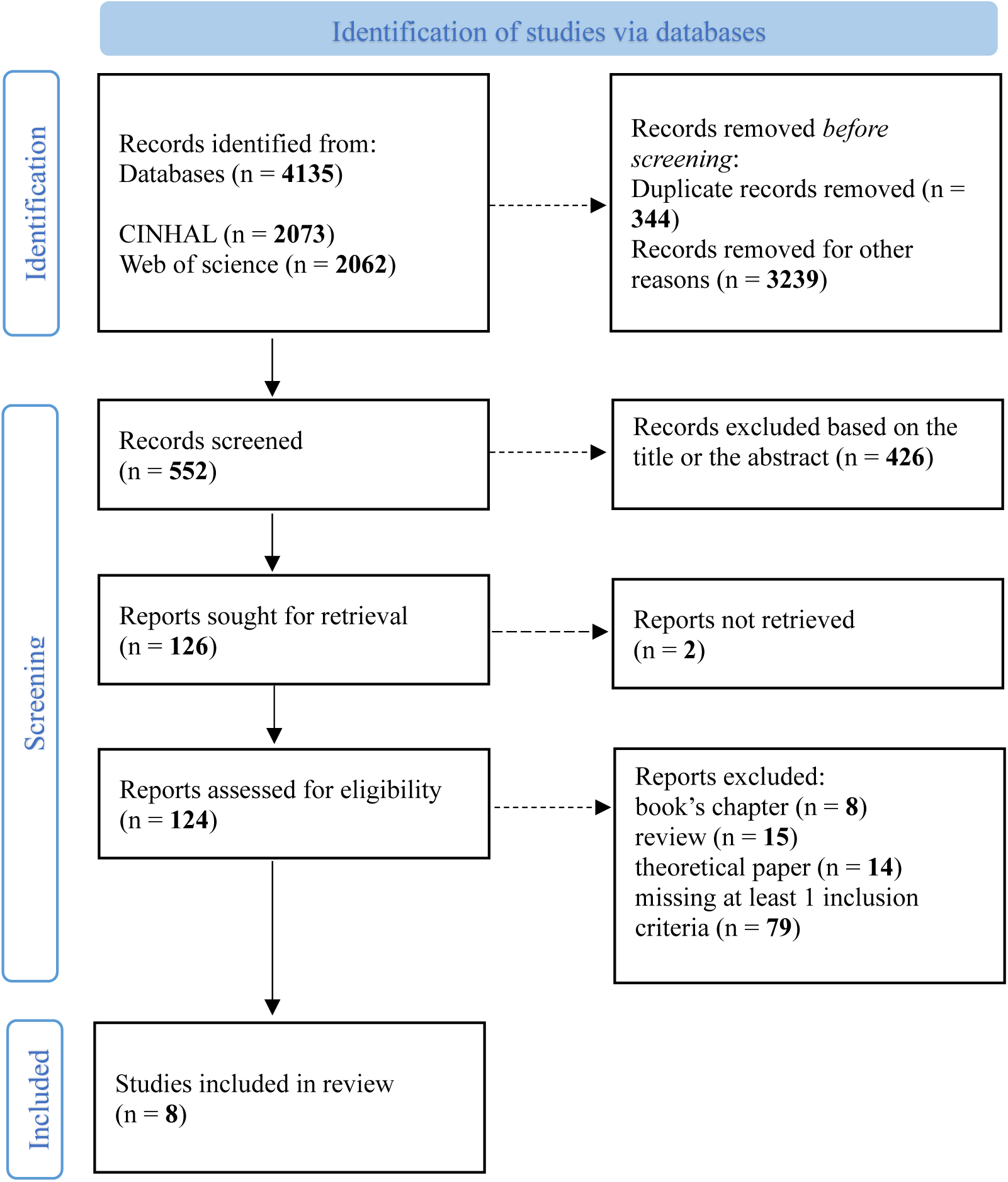
Study selection flow diagram.

### 2.4. Data extraction

The principal data were extracted from selected studies and included the author and the year of publication. Studies featured an experimental design or were intervention studies with an experimental design. The experimental paradigm, the population characteristics, and the assessment and stimuli domains were extracted. We also extracted brain outcome measures, regions of interest, significant results, the conclusion, and potential bias evaluation from the selected studies.

## III. Results

### 3.1. Studies’ design and population characteristics

#### Studies design

All the selected studies had an experimental design, except for one intervention study with a control group (Zhao & Kuhl, 2016). The studies’ main characteristics and a short description of the designs are presented in Table 1.

**Table 1.** Description of experimental design, population, stimuli domain, and aim.

| Author, reference | Experimental design | Population<br>Number, age (mean ± standard deviation; range; sex)<br>Infants' state at test | Stimuli domain<br>Type of stimuli and description | Aim |
| --- | --- | --- | --- | --- |
| 1. Dumont et al. (2022) | Vibrotactile stimulation-omission paradigm: fixed vs. jittered ISI (not predictable stimulus onset). Stimuli were interspersed with omission trials: 1 omission trial every 7-12 vibrations. | 20 preterm neonates (m at birth = $32.1 \pm 0.5$ wGA, m at test = $33.3 \pm 0.2$ corrected wGA; age range at test: 33.0-33.4 corrected wGA; 11 females).<br>Infants were asleep. | Tactile stimuli.<br>Vibration with a palm vibrator, 3 sec. long vibrations interspersed with 5 sec. long intervals. 10 omissions among 84 stimuli. 13 min. total length. | To assess the ability of premature newborns to form a sensory prediction based on a tactile sequence. |
| 2. Edalati et al. (2022) | Auditory rhythmic regularities violations paradigm: oddball paradigm with high-probability standard rhythm ($p = 79\%$) and infrequent deviant rhythm trials ($p = 21\%$). Two control sequences without any standard trial in between. | 20 preterm neonates (m at birth = $31.48 \pm 1.23$ wGA; m at test = $33 \pm 1.44$ wGA; NA; 8 females).<br>Infants were asleep. | Auditory stimuli.<br>Auditory rhythm in 2/4 m, 60 beats/min. Intertone intervals: 1000, 500, 500 ms (standard rhythm), and 1000, 250, 750 ms (deviant rhythm). | To test whether preterm newborns (30-34 wGA) detect temporal regularities in rhythmic patterns.<br>To test the presence of top-down predictions (rhythmic deviations). |
| 3. Háden et al. (2015) | Temporal structure detection paradigm: trains of complex tones varying in pitch (F0) | 30 full-term neonates (m at birth = $39.7 \pm 1$ wGA; age | Auditory stimuli. | To evaluate if the newborn brain is sensitive to the |
|  | across trains. A pitch was randomly selected for each train. Trains consisted of 8–24 tone repetitions (“slow” rate tones in the 1<sup>st</sup> part of the train and “fast” rate tones in the 2<sup>nd</sup> part), followed by a silent interval. | range at test: 1-3 days; 14 females).<br>Infants were asleep. | Trains of complex tones with 8 different pitches taken from the C major scale: C3, D3, E3, F3, G3, A3, B3, and C4.<br>Total duration: 22 min. | overall probability of ascending vs. descending pitch steps in the stimulus block at different repetition rates. |
| 4. Maffongelli et al. (2018) | Structural violations of an observed goal-directed action sequences paradigm: control vs. violation condition (temporal inversion of two adjacent steps of an action sequence). | 14 6-7 months old infants (m = 200 days ± NA; 179-235 days; 7 females).<br>NA | Visual stimuli.<br>Sequences of familiar goal-directed actions are shown as a series of still pictures. | To investigate whether or not a manipulation of the hierarchical organization of an observed familiar action sequence results in differential neurophysiological processing already in infants at the age of 6 to 7 months. |
| 5. Mento and Valenza (2016) | Audio-visual paradigm: oddball paradigm simulating a “Peekaboo” game. Standard (p = 0.77) vs. delayed (p = 0.23) conditions. | 15 9 months old infants (m = 283 days ± 23.9; 252-333 days; 5 females).<br>NA<br>14 adults (m = 25 years ± 3.2; 20-32 years; 9 females). | Visual and auditory stimuli.<br>Visual stimuli: pictures of real female faces, which could be covered by hands or bare-faced and smiling.<br>Auditory stimuli: digitized samples of the notes C<sub>4</sub>, E<sub>4</sub>, G<sub>4</sub>, | To understand whether (1) anticipatory neural mechanisms instantiated by temporal expectancy are already operating early in life and (2) these rely on similar or different spatiotemporal |
|  |  |  | and C<sub>5</sub> played by different musical instruments - i.e., trumpet, oboe, saxophone, and piano. | dynamics as compared to adults. |
| 6. Otte et al. (2013) | Auditory oddball paradigm: violation of temporal regularities. Regular 300 ms ISI between standard tones ($p = 0.7$).<br>3 deviants ($p = 0.1$ each):<br>- 100 ms ISI.<br>- 2 spectral deviants. | 76 2 months old infants ($m = 70.1$ days ± 6.2 days; NA; 46 females).<br>36 waking (20 females) and 40 sleeping (26 females) infants. | Auditory stimuli.<br>Four types of tones: one standard (300 ms ISI) and three deviants. One deviant, identical to the standard, preceded by a 100 ms ISI. Two spectral deviants (white noise sound and novel sound). | To shed light on whether infants can identify violations of temporal regularities (detection of ISI shortenings).<br>To examine the effects of the state of alertness on the MMR. |
| 7. Winkler et al. (2009) | Meter violation paradigm. Standard patterns: base pattern and 3 standard deviants with omission on the lowest metrical level ($p = 22.5\%$ each).<br>1 deviant pattern ($p = 10\%$) with metrically salient omissions in the base pattern.<br>1 control pattern repeating the deviant pattern 100% of the time. | 14 newborn infants (NA; age range at birth: 37-40 wGA, age range at test: 2-3 days; 3 females).<br>Infants were asleep. | Auditory stimuli.<br>Two-measure rock drum accompaniment pattern composed of snare, bass, and hi-hat spanning 8 equally spaced positions. | To investigate whether newborns can extract the beat of the rhythmical sequence (stronger expectation). |
| 8. Zhao and Kuhl (2016) | Intervention study with experimental design. | 39 9-months old infants (NA; NA; NA). Intervention group ($n = 20$) and control group ($n = 19$). | Intervention phase: multimodal, social, and repetitive stimuli (synchronization of | To test whether the intervention enhanced infants' general ability to |
|  | Auditory oddball paradigm (standard, $p = 85\%$; deviant, $p = 15\%$).<br>Effects of a laboratory-controlled music intervention on infants' neural processing of temporal structure in music and speech: music (intervention group) vs. play (control group) activities for 12 sessions. | Inclusion in the analysis of the MEG recordings of 36 participants in the music condition (18 from intervention, 6 females; 18 from control, 9 females) and 35 participants in the speech condition (16 from intervention, 4 females; 19 from control, 10 females).<br>NA | the infants' movements to the musical beats in the intervention group vs. play without music in the control group).<br>Testing phase: auditory stimuli. Music condition: triple meter structure (strong complex tone combined with two weak complex tones with sound-onset-asynchrony of 300 ms). Speech condition: disyllabic nonword speech (combination of synthesized syllables with silent gaps in between). | extract temporal structure and generate more robust predictions about future stimuli in complex auditory sounds. |

More specifically, four studies used auditory rhythmic meter, temporal regularities, or goal-directed action structural violations paradigms (Edalati et al., 2022; Maffongelli et al., 2018; Otte et al., 2013; Winkler et al., 2009). Two studies used an auditory or audio-visual paradigm (Mento & Valenza, 2016; Zhao & Kuhl, 2016). The last two studies used a vibrotactile stimulation-omission paradigm (Dumont et al., 2022) and a temporal structure detection paradigm (Háden et al., 2015).

#### Population

The selected studies included a total population of 228 infants. Among these infants, 40 were preterm (32-33 wGA at birth and 33 wGA at test) neonates (Dumont et al., 2022; Edalati et al., 2022). Two studies were conducted on a total of 44 full-term (39-40 wGA at birth and 1-3 days at test) newborns (Háden et al., 2015; Winkler et al., 2009). One study was conducted on 76 2-month-old participants (Otte et al., 2013). Another study was conducted on 14 6- to 7-month-old infants (Maffongelli et al., 2018). The last two studies recruited 9-month-old infants, for a total of 54 infants (Mento & Valenza, 2016; Zhao & Kuhl, 2016).

In four studies, infants were asleep when tested (Dumont et al., 2022; Edalati et al., 2022; Háden et al., 2015; Winkler et al., 2009). In the study by Otte et al. (2013), 36 infants were awake, and 40 were asleep at the time of testing. Nothing has been mentioned about the infant’s state at the moment of the test in the other studies.

One selected study (Mento & Valenza, 2016) has also been conducted on 14 adults (adults’ results have not been included in this systematic review but are presented in the supplementary material; cf. Figures 4 and 5).

### 3.2. Stimuli domains

Five of eight experimental paradigms used auditory stimuli like auditory rhythm, trains of complex tones with different pitches, different types of tones, a two-measure rock drum, or a triple meter structure (Edalati et al., 2022; Háden et al., 2015; Otte et al., 2013; Winkler et al., 2009; Zhao & Kuhl, 2016).

In one study, audio and visual stimuli were paired, where Mento and Valenza (2016) used pictures of real female faces along with digitized samples of notes played by different musical instruments. One study used visual stimuli consisting of sequences of familiar goal-directed actions (Maffongelli et al., 2018). Finally, tactile stimuli have been used only in one study (Dumont et al., 2022).

A more detailed description of the stimuli used in each study is available in Table 1.

### 3.3. Aims

The declared aim of three studies was to assess infants’ abilities to extract temporal regularities in a series of stimuli to form sensory predictions (Dumont et al., 2022; Edalati et al., 2022; Zhao & Kuhl, 2016). Two studies focused on infants’ ability to develop temporal expectations, with a particular interest in anticipatory neuronal mechanisms and their similarity to adults’ spatiotemporal dynamics (Mento & Valenza, 2016; Winkler et al., 2009). The goal of the study by Otte et al. (2013) was to understand whether infants could identify violations in temporal regularities. The last two studies aimed to evaluate infants’ brain sensitivity to the manipulation of pitch steps at different repetition rates (Háden et al., 2015) and the impact of the manipulation of hierarchical organization on neurological processing (Maffongelli et al., 2018).

### 3.4. Brain outcome measures and regions of interest

#### Brain outcome measures

Of the eight selected studies, six used electroencephalography (EEG) as a brain measure. More information about the type of EEG used or the number of channels can be found in Table 2. The measured brain responses were ERPs (e.g., CNV, MMN/MMR; Edalati et al., 2022; Mento & Valenza, 2016; Otte et al., 2013; Winkler et al., 2009). The last two studies used Diffuse Correlation Spectroscopy (or DCS) (Dumont et al., 2022) and magnetoencephalography (or MEG) (Zhao & Kuhl, 2016) as brain measures.

**Table 2.**
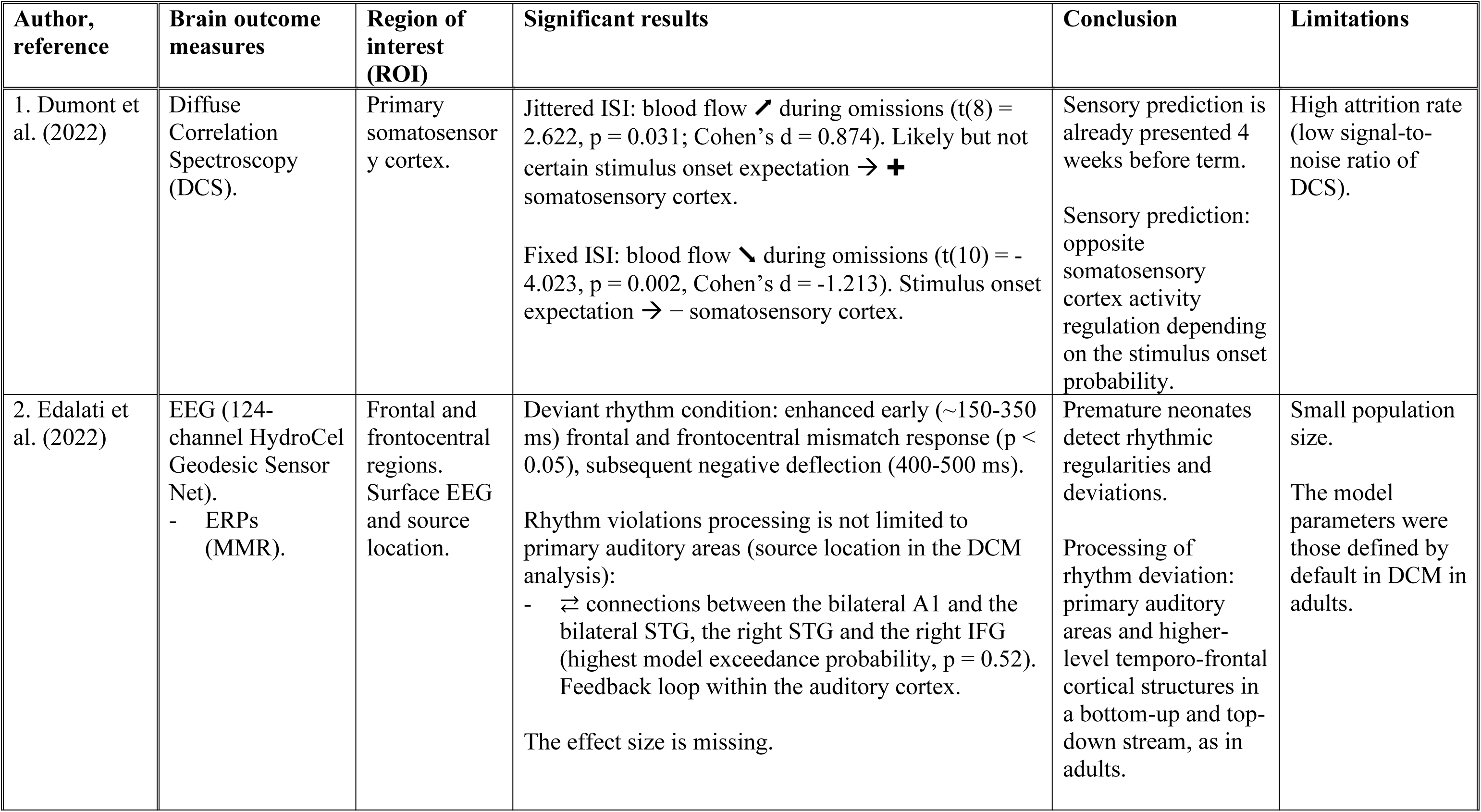
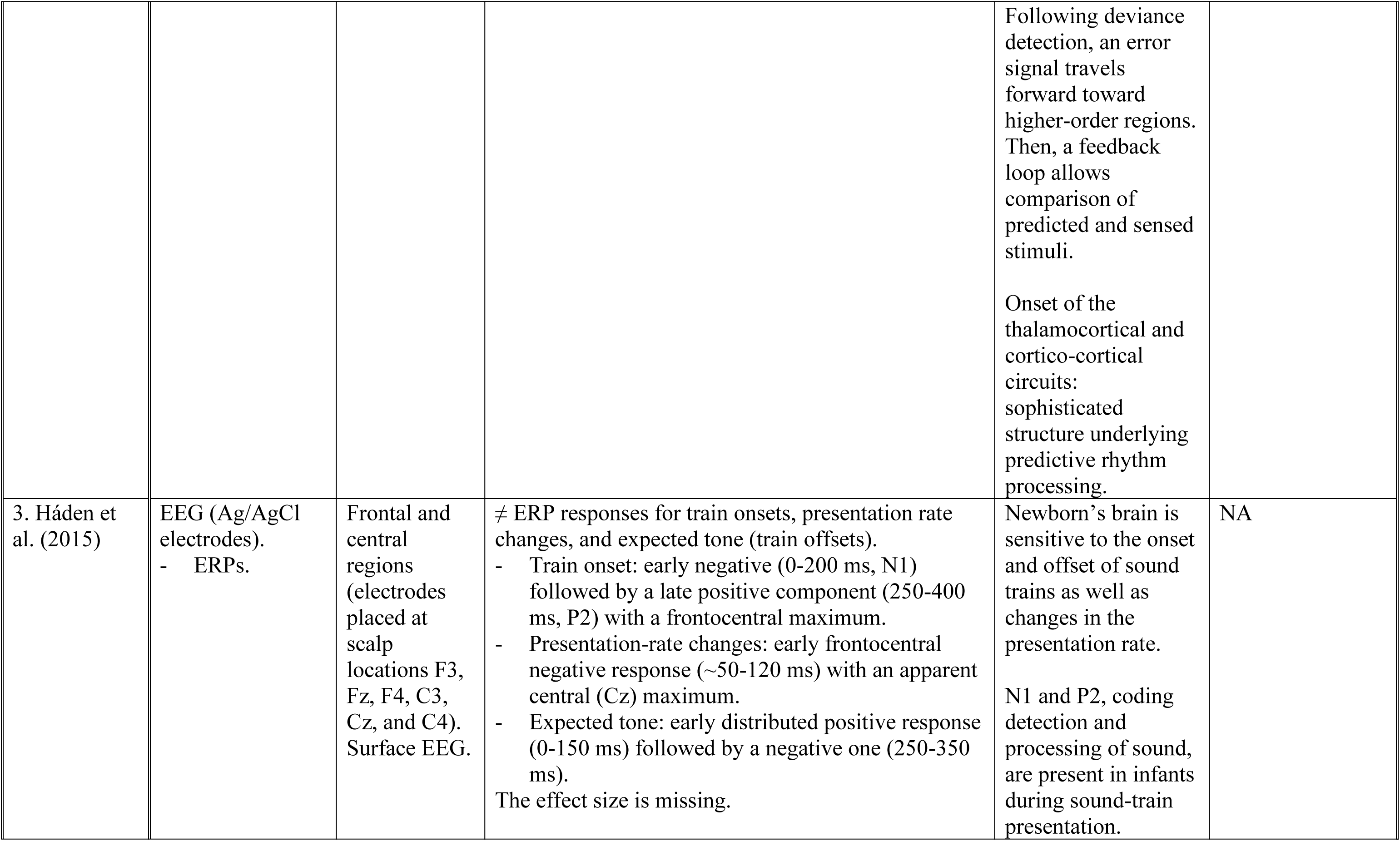
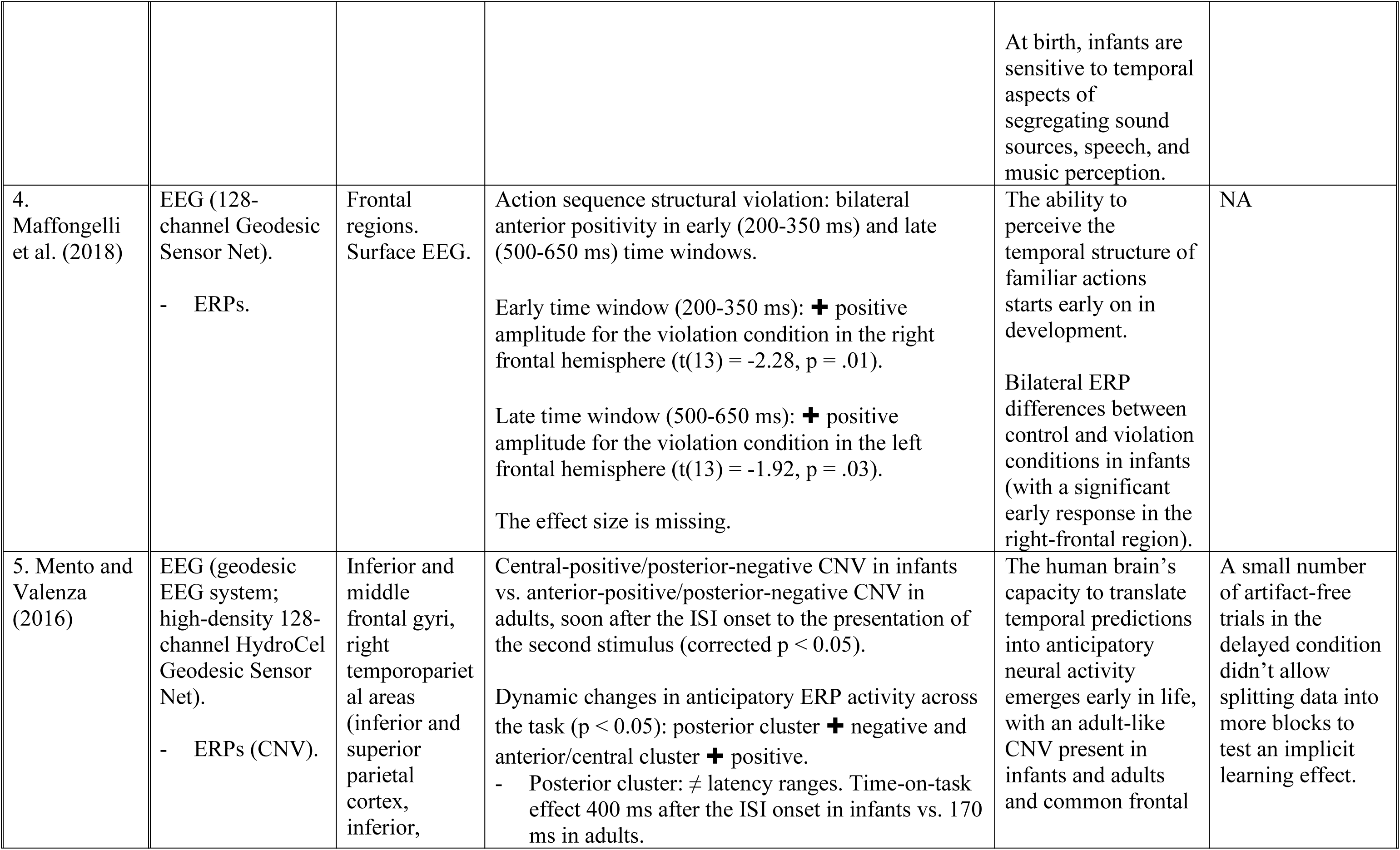
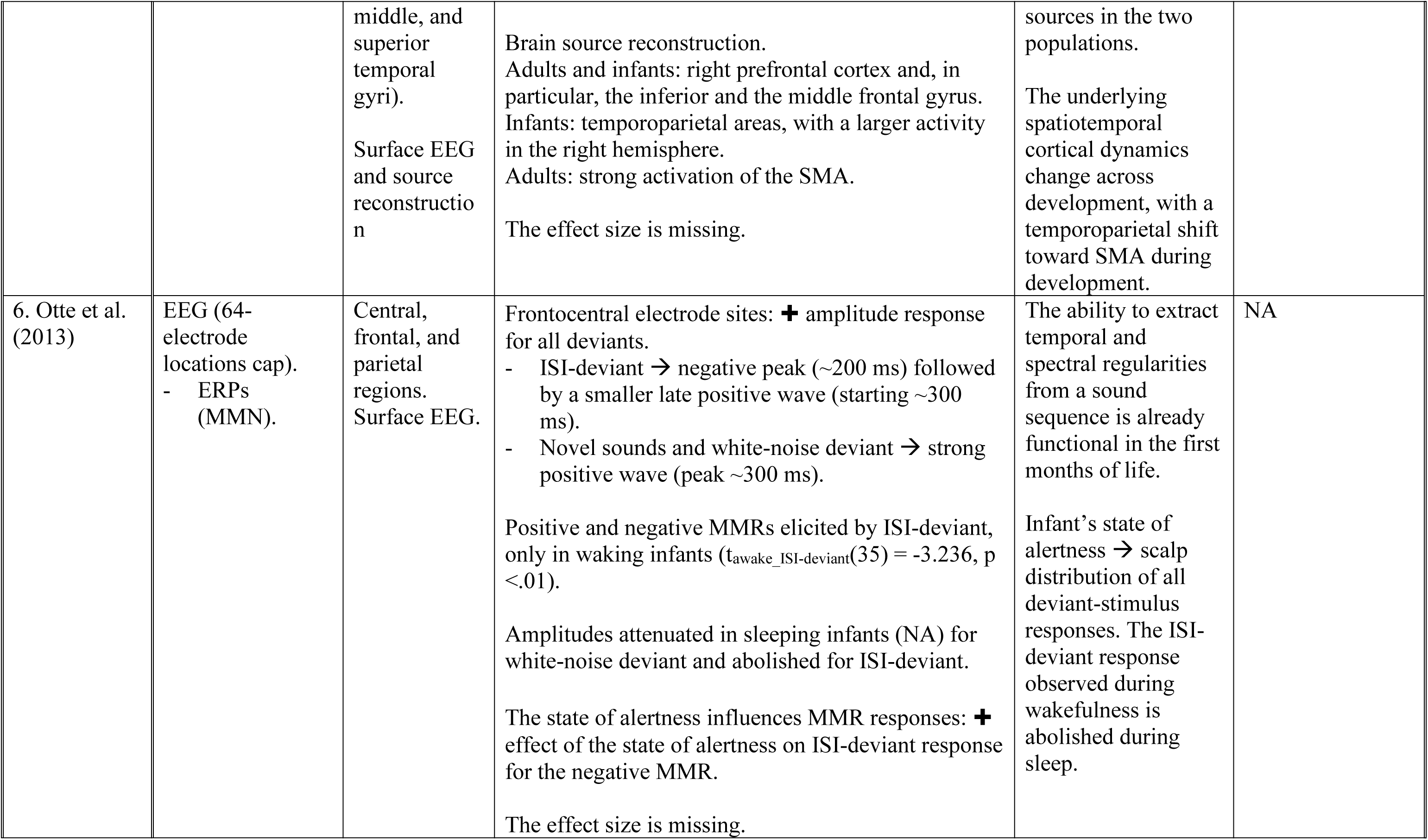
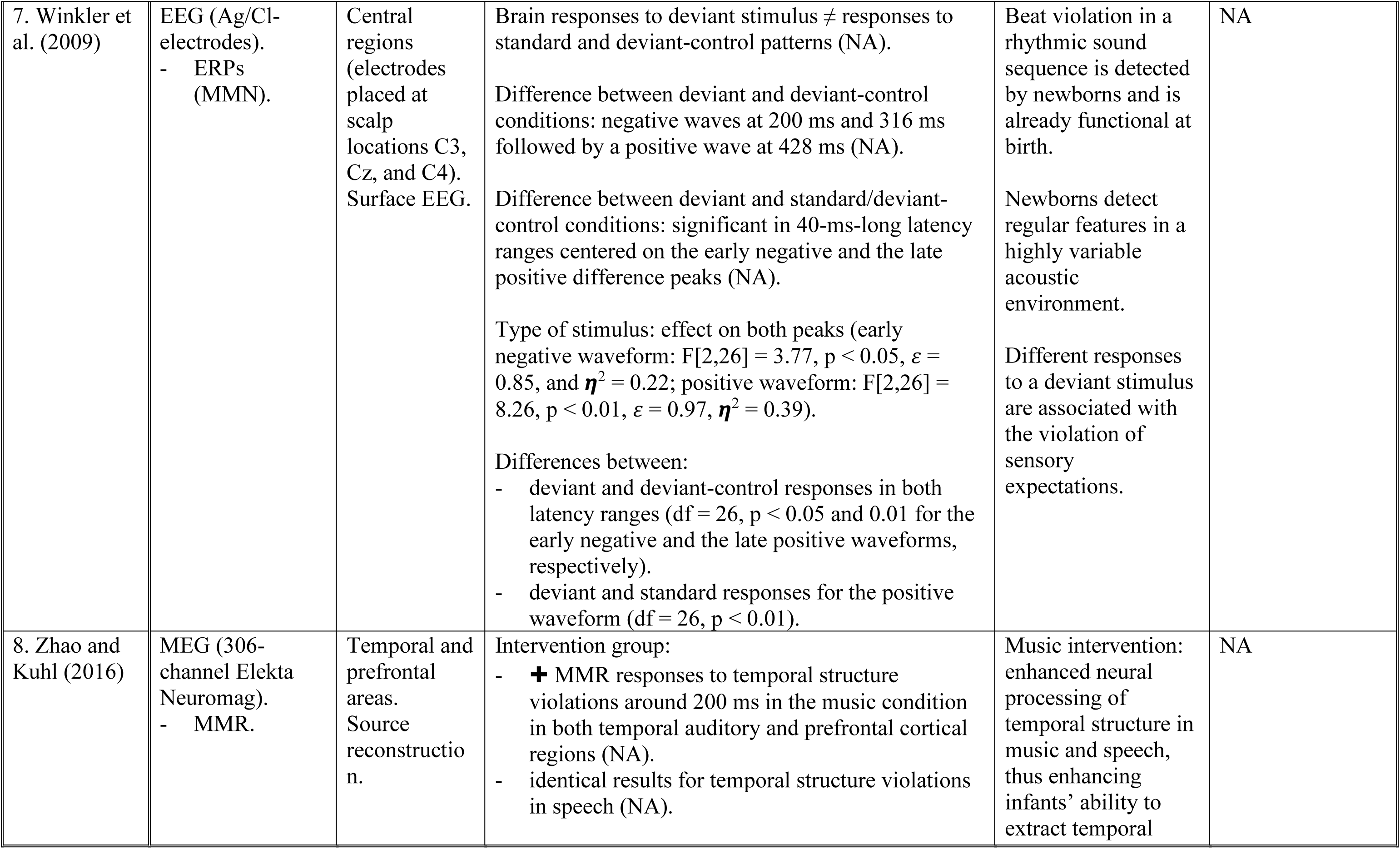
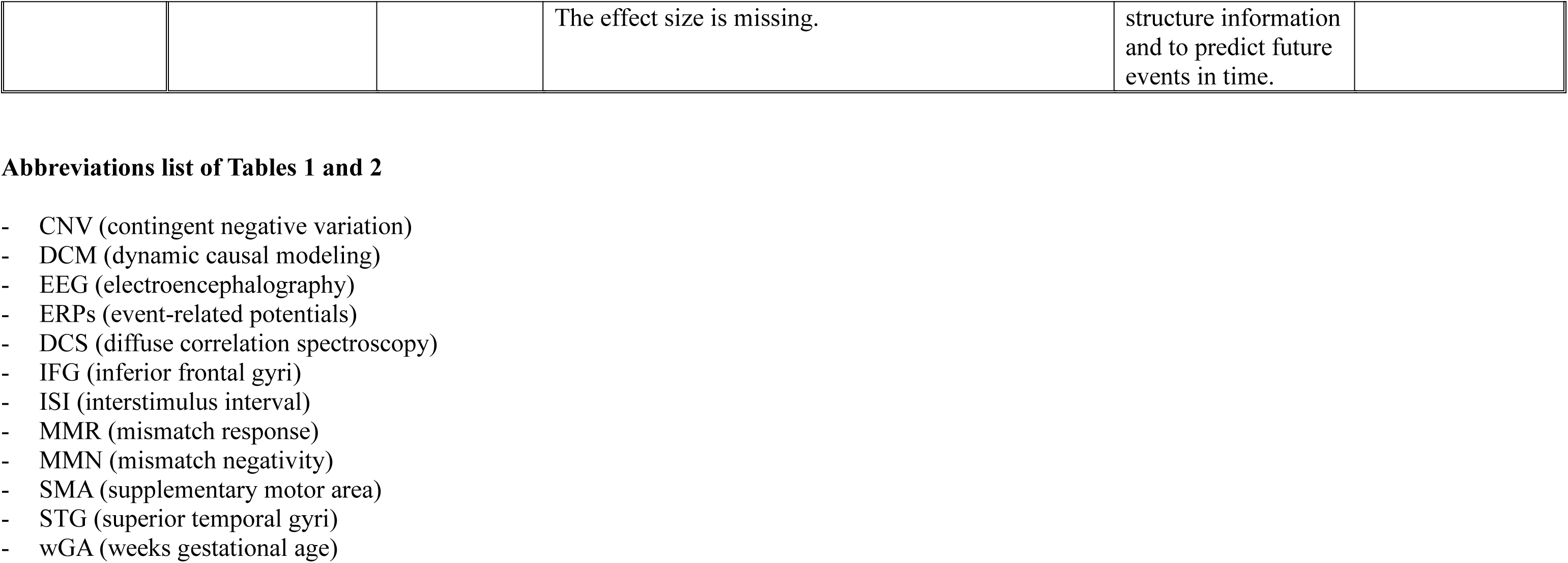
Description of brain outcome measures, region of interest, significant results, conclusion, and limitations.

#### Regions of interest

Three studies focused on one specific brain region (Dumont et al., 2022; Maffongelli et al., 2018; Winkler et al., 2009), whereas other studies investigated multiple brain regions. Most of the selected studies (5/8) concentrated on specific brain activations for temporal prediction in the frontal regions (Edalati et al., 2022; Háden et al., 2015; Maffongelli et al., 2018; Mento & Valenza, 2016; Otte et al., 2013) and three studies in central areas (Háden et al., 2015; Otte et al., 2013; Winkler et al., 2009).

Three studies focused on the involvement of parietal (Otte et al., 2013), temporal (Zhao & Kuhl, 2016), and temporoparietal regions (Mento & Valenza, 2016) in temporal prediction.

One study focused on specific brain activations for temporal prediction in frontocentral regions (Edalati et al., 2022), and another one in prefrontal areas (Zhao & Kuhl, 2016). The last study involved the primary somatosensory cortex (Dumont et al., 2022).

It’s important to note that six out of eight studies (Edalati et al., 2022; Háden et al., 2015; Maffongelli et al., 2018; Mento & Valenza, 2016; Otte et al., 2013; Winkler et al., 2009) used surface EEG, while Zhao and Kuhl (2016) used source reconstruction in a MEG paradigm.

Two studies (Edalati et al., 2022; Mento & Valenza, 2016) used surface EEG and brain source location or reconstruction. Indeed, in their studies, Edalati et al. (2022) used DCM analysis for brain source location, while Mento and Valenza (2016) extracted the regions of interest (i.e., inferior and middle frontal gyri and the right temporoparietal areas) from a brain source reconstruction.

More detailed information about brain regions of interest is summarized in Table 2.

### 3.5. Results and conclusions of the included studies

Results summarizing the main findings of the included studies will be presented in a potential longitudinal perspective (which has to be nuanced, cf. limitations of the present review), with the infants’ age groups in Figure 2, as well as results extracted from source location and reconstruction in Figure 3.

**Figure 2.**
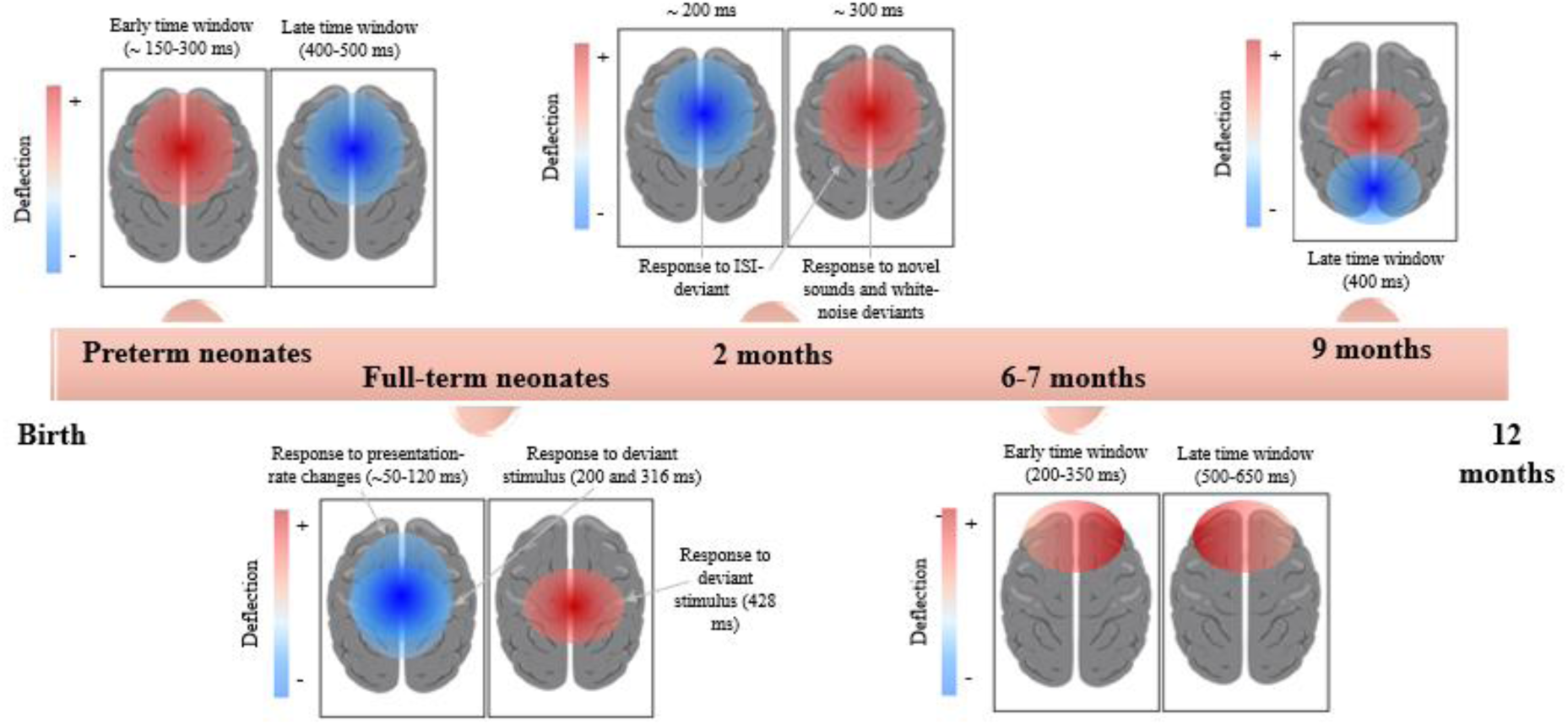
Neural signatures of temporal prediction in infancy at the scalp surface (excerpt from Edalati et al., 2022; Háden et al., 2015; Maffongelli et al., 2018; Mento & Valenza, 2016; Otte et al., 2013; Winkler et al., 2009).

**Figure 3.**
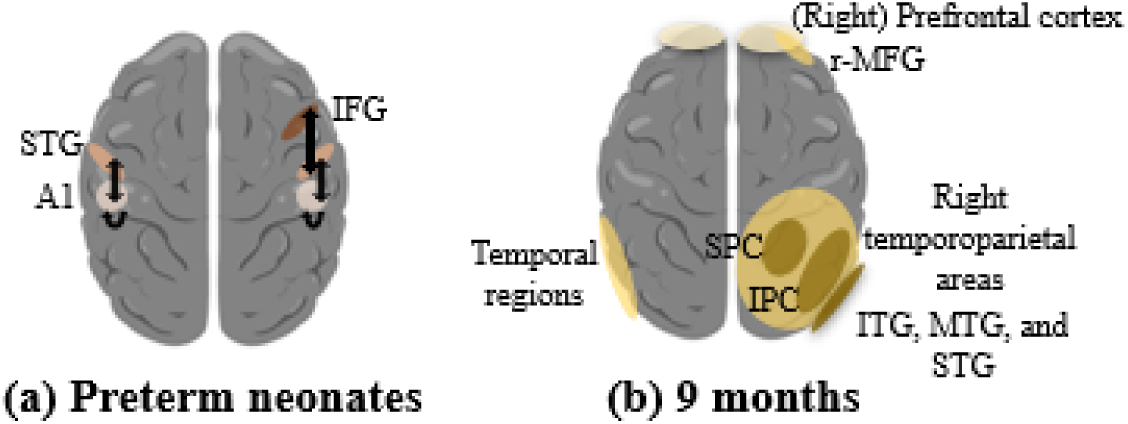
Source location and reconstruction of temporal prediction in (a) preterm neonates and (b) 9-month-old infants (excerpt respectively from Edalati et al., 2022; Mento & Valenza, 2016; Zhao & Kuhl, 2016). (a) In preterm neonates, DCM analysis identified bidirectional connections between the bilateral auditory cortex (A1) and the bilateral superior temporal gyri (STG), the right STG and the right inferior frontal gyrus (IFG), and a feedback loop within the auditory cortex. (b) In 9-month-old infants, source reconstruction identified a cortical activity in the right prefrontal cortex, including the inferior and the middle frontal gyrus (r-MFG) and in the right temporoparietal areas (with an extending activity over the inferior (IPC) and the superior (SPC) parietal cortices, the inferior (ITG), the middle (MTG), and the superior (STG) temporal gyri). Moreover, in 9-month-old infants, a MEG study identified brain activity in the temporal (auditory) and prefrontal cortices.

In preterm neonates aged 33 (corrected) weeks of gestational age, one study showed that in a vibrotactile paradigm, omissions in the jittered interstimulus interval (or ISI) condition (i.e., in likely but not certain stimulus onset expectation) induce an activation of the somatosensory cortex, whereas a deactivation is observed in this region during omission when the stimulus onset was expected (in the fixed interstimulus interval condition) (Dumont et al., 2022). In preterm babies, another study (Edalati et al., 2022) has shown that in an auditory rhythmic regularities violation paradigm, a deviant condition elicited an early (∼150–350 ms) frontal and frontocentral mismatch response (MMR), followed by a negative deflection (400-500 ms). More specifically, using Dynamic Causal Modeling (or DCM) for source location, authors showed that MMR is not limited to the primary auditory cortex but consists of a feedback loop within the auditory cortex, associated with bidirectional connections between the bilateral primary auditory cortex and the bilateral superior temporal gyrus, the right superior temporal gyrus and the right inferior frontal gyrus (Edalati et al., 2022).

In infants born prematurely, sensory prediction following a vibrotactile stimulation is already present and implies the primary somatosensory cortex with an opposite activity regulation depending on the stimulus onset probability (Dumont et al., 2022). Likewise, the infant’s brain extracts rhythmic regularities and detects deviations from them. This operation, as with adults, also relies on higher-level temporo-frontal brain structures in a bottom-up and top-down fashion (Edalati et al., 2022). Indeed, after deviance detection, an error signal is sent toward higher-order regions, followed by a feedback loop that enables the comparison between predicted and sensed stimuli (Edalati et al., 2022).

At birth, a full-term infant’s brain is already sensitive to changes in the rate at which sounds occur, as shown by the elicited early frontocentral negative response (∼50-120 ms) (Háden et al., 2015) and presents a different response to a deviant stimulus (Winkler et al., 2009). Indeed, Winkler et al. (2009) have observed two negative waves (peaking at 200 ms and 316 ms) followed by a positive wave (peaking at 428 ms) in central regions for the difference between the deviant and the deviant control conditions; further post hoc analysis also revealed significant differences between the deviant and the standard response for the positive waveform in the same brain region.

As shown by these studies involving frontal and central regions, at birth, the infant brain is already sensitive to the temporal aspects of sounds and their segregation. It can detect regularities in a highly variable acoustic environment, as well as beat violations in a rhythmic sequence that conflict with infants’ sensory expectations (Háden et al., 2015; Winkler et al., 2009).

In 2-month-old infants, an ISI-deviant elicited a negative peak (∼200 ms) followed by a smaller late positive wave (∼300 ms) in the frontal, central, and parietal regions; whereas novel sounds and white-noise deviants elicited a strong positive wave in the same brain regions (Otte et al., 2013). However, the highest amplitude response has been observed on frontocentral sites for all deviants, although these amplitudes attenuate for white-noise deviants and abolish for ISI-deviants in sleeping infants (Otte et al., 2013). Interestingly, Otte et al. (2013) reported that ISI-deviant elicited positive and negative MMRs only in waking infants, showing an effect of the infant’s state of alertness (awake *vs.* asleep) on the scalp distributions of the MMR responses, with a larger effect of the state of alertness on ISI-deviant response for the negative-going MMR.

Early in life, the infant’s brain can extract temporal regularities from a sound sequence. As shown by the authors, the infant’s state of alertness impacts the fronto-centro-parietal response distribution to deviant conditions (i.e., the response to ISI-deviant observed in waking infants is abolished during sleep) (Otte et al., 2013).

At 6 to 7 months of age, a structural violation in an action sequence elicited a bilateral anterior positivity in early (200-350 ms) and late (500-650 ms) time windows, with a more positive amplitude for the violation condition in the right and left frontal hemispheres, respectively (Maffongelli et al., 2018).

As shown by this study, the temporal structure of familiar goal-directed actions and their violations can be detected in the brains of 6- to 7-month-old infants, resulting in bilateral ERP modifications in the frontal regions (Maffongelli et al., 2018).

At 9 months old, Mento and Valenza (2016) reported anticipatory brain activity, with dynamic changes in anticipatory ERP activity along the task (i.e., the posterior cluster at the surface of the scalp became more negative and the anterior/central cluster more positive indicating, using brain source reconstruction, a larger neural activity in the inferior and middle frontal gyri, and the right temporoparietal areas as the temporal contingency between the two stimuli was implicitly processed by infants’ brain in the delayed condition) in an oddball paradigm simulating a “Peekaboo” game. A laboratory-controlled music intervention (Zhao & Kuhl, 2016) revealed in both temporal auditory and prefrontal cortical regions (using source reconstruction) larger MMR responses (∼200 ms) to temporal structure violations in music and speech in infants of the same age.

These results show that the infant’s brain is already able to translate predictions of temporal time sequences into prefrontal and temporoparietal anticipatory brain activities at 9 months of age, which translates into an adult-like CNV (Mento & Valenza, 2016). This neural processing might be enhanced by a music intervention, as shown by Zhao and Kuhl (2016), for a better prediction of upcoming events.

More detailed information on the studies’ results is summarized in Table 2.

### 3.6. Bias evaluation

#### By the authors of their experimental paradigm

Five studies did not report any bias evaluation. Dumont et al. (2022) reported a high attrition rate. Edalati et al. (2022) reported a small population size (with 20 preterm neonates in total) and used model parameters based on adults. Mento and Valenza (2016) couldn’t test an implicit learning effect due to a small number of artifact-free trials.

## IV. Discussion

The primary objective of this systematic review was to identify the neural signatures of temporal prediction in infancy and to point out the brain signatures of prediction mechanisms, examining their consistency and evolution over time from birth to 12 months of age.

Over a third of the recruited infants were newborns, demonstrating that age gaps in the existing literature enable the exploration of the emergence of temporal predictive abilities. Within the chosen studies, both premature and full-term infants underwent testing at birth, as well as infants aged 2, 6-7, and 9 months. Notably, there is a testing gap between the ages of 2 and 6-7 months and after 9 months, as no studies have been conducted on infants at the age of 12 months. The consistency in measurements allows us to adopt a potential longitudinal perspective in our discussion, spanning from birth to 9 months of age (which has to be nuanced, cf. limitations of the present review).

Regarding the methods of investigation, electroencephalography (EEG) has been largely used in the reported studies, thereby establishing EEG as the gold standard for assessing the brain’s temporal signature of prediction early in infancy. EEG use as a brain measure proves particularly suitable for evaluating temporal and rhythmic brain mechanisms in infants, rather than other brain measurement techniques (i.e., current studies on auditory processing in infancy use EEG as a brain measure; see Edalati et al., 2023, for an example).

Within the chosen studies, two deviations from the EEG used have been noted, with DCS (Dumont et al., 2022) in preterm neonates and magnetoencephalography employed in the assessment of older infants, as indicated by the research conducted by Zhao and Kuhl (2016).

The current systematic review allowed us to draw some conclusions about the localization, temporality, and polarity (at the surface of the scalp) of the neural signatures of temporal prediction in infants aged 0 to 9 months, as well as their potential longitudinal development over time (which has to be nuanced, cf. limitations of the present review).

Infants’ temporal predictions, based on surface location, appear to primarily focus on the brain’s centro-anterior regions, including the frontal and central areas.

In preterm, full-term, and 2-month-old infants, bilateral ERP modifications at a similar timing have been observed at the surface of the scalp in fronto-central areas in response to rhythm violation or deviance (Edalati et al., 2022; Háden et al., 2015; Otte et al., 2013; Winkler et al., 2009). These ERP modifications are characterized by an early negative deflection followed by a late positive wave in full-term neonates and 2-month-old infants (Háden et al., 2015; Otte et al., 2013; Winkler et al., 2009); whereas an opposite polarity change is observed in preterm neonates (Edalati et al., 2022). Regarding the temporality of these ERPs in full-term and 2-month-old infants, a common 200-ms negative wave is observed in both paradigms (Otte et al., 2013; Winkler et al., 2009); however, the late positive deflection appears earlier in a 2-month-old infant’s brain (Otte et al., 2013). This earlier response to deviance might indicate, if similar brain processes are involved, a differentiation of deviance detection mechanisms only a few months after birth. The observed polarity and temporality differences between preterm, full-term, and 2-month-old infants (Edalati et al., 2022; Háden et al., 2015; Otte et al., 2013; Winkler et al., 2009) might be explained by the use of different experimental paradigms in each study, as well as different brain mechanisms and/or network implications, and their neural sources or desynchronization/synchronization.

At 6 to 7 months old, bilateral frontal positivity, both in early and late time windows, is observed in response to a structural violation in action sequences (Maffongelli et al., 2018). The difference in observed ERP polarity (in comparison to the above-mentioned studies) could be explained by the nature of the task, which involves action sequences.

Moreover, throughout infants’ development, a noticeable shift in activated areas might be observed (measured at the surface of the scalp): central and frontal activity remains highly specific to this early cognitive ability, as observed in studies conducted in 6-7-month- and 9-month-old infants (Maffongelli et al., 2018; Mento & Valenza, 2016), but may extend to posterior areas as they develop (Mento & Valenza, 2016).

Some exceptions in the reported pool of activated brain areas have been noticed (Dumont et al., 2022). This variation can be attributed to the specific paradigm in this DCS study, involving vibrotactile stimulation that elicited distinct somatosensory responses. It is worth noting that we do not posit the primary somatosensory cortex as exclusively dedicated to predictions. Despite the divergence in paradigm design from other studies, this form of stimulation is particularly intriguing and judicious, given that newborns’ sensory perception is fundamentally intermodal.

The inquiry into the potential lateralization of prediction mechanisms in infants’ brain signatures is interesting. However, the selected studies lack sufficient information to address this question comprehensively, with only two studies providing specificity on lateralization, based on source location and reconstruction, particularly focusing on the right prefrontal cortex and the right temporoparietal regions (Mento & Valenza, 2016), as well as the right STG and right IFG during rhythm violations processing in an auditory paradigm (Edalati et al., 2022).

In one included study (Mento & Valenza, 2016), adults were also tested using the same paradigm as in infants; based on this article, figures presenting the neural signatures at the scalp surface and the source reconstruction of temporal prediction in adults can be found in the supplementary material (cf. Figures 4 and 5).

Some limitations of the included studies emerged, though they did not diminish the significance of their findings. In our view, a term-born control population was missing in Dumont et al. (2022) and Edalati et al. (2022) as a follow-up measure in the first study.

Maffongelli et al. (2018) had a limited sample size. In Dumont et al. (2022), brain measurements were limited to the specified region of interest, and we were unable to encompass the chosen brain regions to gain a better understanding of whole-brain signatures of prediction. Edalati et al. (2022) employed dynamic causal modeling with adults’ model parameters to draw inferences regarding the neuronal architecture underlying the observed mismatch responses in preterm infants.

Moreover, in numerous studies, the infants’ state during testing was not specified, posing a limitation to the corpus and, consequently, our conclusions. Studying very young infants is inherently challenging due to their limited moments of wakefulness; however, as they age, the frequency of wakeful periods increases, providing more data on alert infants. The absence of consistent reporting on infants’ states during testing or the variability in infants’ alertness levels during testing can pose challenges. As noted by Otte et al. (2013), the state of alertness in infants significantly influences brain responses, with sleeping infants exhibiting attenuated amplitudes and only waking infants displaying an ISI-deviant response.

Finally, there is a terminological limitation in the studies included. Indeed, the terms “anticipation”, “expectation”, “prediction”, or “violation” are often used without being clearly defined, leading to terminological and, therefore, theoretical confusion.

### 4.1 Limitations of the present review and recommendations for further investigations

As a first step towards synthesizing the existing literature on the brain mechanisms of temporal prediction in early ages, this systematic review presents certain limitations. First, the focus only on the temporal aspects of prediction (as a subcategory) as a methodological choice of the present review, inevitably restricted the number of articles included, which could potentially limit the impact and generalizability of the present results to all, and to some extent, complex and studied using a variety of experimental paradigms, prediction mechanisms. Focusing on the temporal dimension of predictions may offer valuable insights into the core mechanisms underlying certain atypical trajectories – such as those observed in preterm infants, who often exhibit early difficulties in temporal sensory integration and interpersonal synchrony (Feldman, 2007a; Lester et al., 1985). Second, considering the structural limitations of the current field, available studies on the topic are scattered, resulting in a limited number of subjects examined across an extensive age range, and are understandably heterogeneous. This is due to inherent challenges in studying early developmental stages in infants, as well as considerable variations in experimental paradigms used and selected brain regions of interest. In this context, the potential developmental perspective suggested in this review must be interpreted with caution, although predictive abilities are present from the very beginning of life. Third, a comparison of basal EEG activity measures at the scalp surface (i.e., the morphological characteristics of scalp-recorded waveforms) or brain source location/reconstruction for the concerned studies (Edalati et al., 2022; Mento & Valenza, 2016; Zhao & Kuhl, 2016) from birth to 9 months of age, is challenging and has to be interpreted with caution. Indeed, the recorded brain activity and the subsequent source location/reconstruction and its comparison across age may be influence by both maturational (e.g., head size and shape, open fontanelle in neonates, scalp thickness, brain maturation, structural differences such as connectome changes, brain volume conduction, or dipole orientation; Collin & Van Den Heuvel, 2013; Paus, 2024; Reynolds & Richards, 2009) and methodological factors such as cap montage (especially the reference used; Rosenfeld, 2000). For instance, fixed references (as linked mastoids used in low-density EEG) redistribute electrical activity differently from the average reference commonly used with high-density systems (Chella et al., 2016; Rosenfeld, 2000).

However, these last two limitations (i.e., the resulting difficulty in reliably comparing scalp distribution and waveform polarity across age groups due to the variety of experimental tasks used) could be nuanced by the use of source-level brain analyses, providing significant indicators of the neural source underlying scalp activity, despite challenges related to forward modelling and age-specific head models (Michel & Brunet, 2019). In this context, network analysis in preterm neonates from Edalati et al. (2022), as well as the study conduct in 9-months old infants by Mento and Valenza (2016) might be particularly relevant as it report a widespread (pre)fronto-temporo-parietal activation, potentially required the integration of posterior regions (engaged in perceptual processing) with anterior regions (linked to domain general functions) in upcoming studies.

Regarding the conclusion of the present review, as well as the current state of the field, we recommend including a typically developing population in further studies of preterm neonates. Indeed, this could enable a more comprehensive comparison and interpretation of the observed effects, providing valuable insights into the developmental trajectory of the studied phenomena and enhancing the generalizability of the findings. Moreover, we recommend meticulously assessing and ensuring uniformity in infants’ states during testing to improve the reliability of findings, as variations in behavioral state have been shown to influence neural responses significantly (Otte et al., 2013). Further investigations would also need to confirm the potential effect of music exposure or rhythmical joint activities on neural integration across sensory and higher-order brain regions observed by Zhao and Kuhl (2016) in our corpus. Finally, potentially similar brain processes of prediction and their differentiation through the infant’s first year of life would be interesting to investigate longitudinally in further studies, using a unique temporal deviance detection experimental high-density EEG paradigm in different age groups. This perspective enables temporal prediction to be linked to the development of time perception in infants, for instance. In speech processing and production, rhythm is closely associated with speech motor cortex development (Poeppel & Assaneo, 2020), shaping the evolution of articulation and rhythm in infant-directed speech. Adults adapt their speech speed to the infant’s perceptual and temporal abilities: from slow and regular to faster and more variable rates (Kitamura & Burnham, 2003; Rosslund et al., 2024). These temporal adjustments help infants acquire predictive models of speech – initially by minimizing prediction errors and later by increasing lexical density, facilitating further vocabulary and syntax acquisition (Cristia, 2013; Weisleder & Fernald, 2013).

## V. Conclusion

Emerging even before birth, newborns demonstrate temporal predictive abilities within a structured temporal framework, with EEG serving as the gold standard for studying these early cognitive processes. While earlier research relying on behavioral observations suggested that predictive abilities are present from birth, neurophysiological investigations have since confirmed and deepened our understanding of these early-emerging capacities. Although these abilities have a broad cerebral localization, the majority of studies show that they are concentrated in the anterior and medial parts of the brain, particularly in the frontal and central areas, with consistency across ages. Temporal prediction mechanisms appear to transition from early sensory-driven responses to more complex, top-down predictive processing over the first year of life. The development of these mechanisms relies on both biologically driven and experience-dependent processes, with external factors such as shared rhythmic and musical activities potentially shaping predictive abilities, as reflected in neural responses. This underscores the dynamic nature of temporal prediction development and highlights the integration of sensory and higher-order brain regions as infants mature.

