## Supplementary material for "The origins of time: a systematic review of the neural signatures of temporal prediction in infancy"

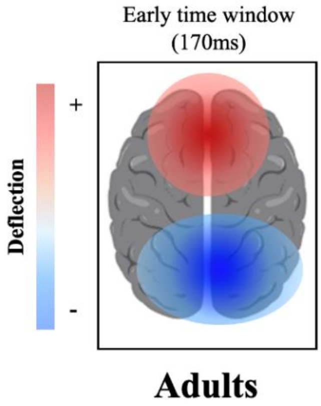


***Figure 4*. Neural signatures of temporal prediction in adults at the scalp surface (excerpt from Mento & Valenza, 2016)**

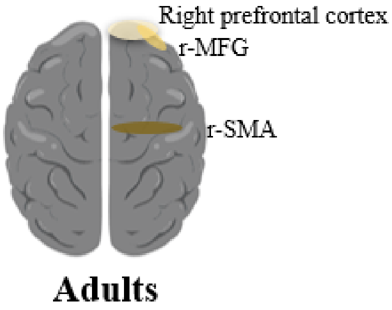


***Figure 5*. Source reconstruction of temporal prediction in adults (excerpt from Mento & Valenza, 2016).** In adults, source reconstruction identified cortical activity in the right prefrontal cortex, including the inferior and the middle frontal gyrus (r-MFG), and in the right supplementary motor area (r-SMA).
